# Spatio-Temporal Gene Discovery For Autism Spectrum Disorder

**DOI:** 10.1101/256693

**Authors:** Utku Norman, A. Ercument Cicek

## Abstract

Whole exome sequencing (WES) studies for Autism Spectrum Disorder (ASD) could identify only around six dozen risk genes to date because the genetic architecture of the disorder is highly complex. To speed the gene discovery process up, a few network-based ASD gene discovery algorithms were proposed. Although these methods use static gene interaction networks, functional clustering of genes is bound to evolve during neurodevelopment and disruptions are likely to have a cascading effect on the future associations. Thus, approaches that disregard the dynamic nature of neurodevelopment are limited in power. Here, we present a spatio-temporal gene discovery algorithm for ASD, which leverages information from evolving gene coexpression networks of neurodevelopment. The algorithm solves a variant of prize-collecting Steiner forest-based problem on coexpression networks to model neurodevelopment and transfer information from precursor neurodevelopmental windows. The decisions made by the algorithm can be traced back, adding interpretability to the results. We apply the algorithm on WES data of ^3^,^871^ samples and identify risk clusters using BrainSpan coexpression networks of early- and mid-fetal periods. On an independent dataset, we show that incorporation of the temporal dimension increases the prediction power: Predicted clusters are hit more and show higher enrichment in ASD-related functions compared to the state-of-the-art. Code is available at http://ciceklab.cs.bilkent.edu.tr/ST-Steiner/.

## I. INTRODUCTION

Autism Spectrum Disorder (ASD) is a common neurodevelopmental disorder that affects *∼*1.5% of the children in the US (Autism and Investigators, 2014). Recent whole exome sequencing (WES) efforts have paved the way for identification of dozens of ASD risk genes (O’Roak *et al.*, 2012; Iossifov *et al.*, 2012; Neale *et al.*, 2012; Sanders *et al.*, 2012; De Rubeis *et al.*, 2014; Iossifov *et al.*, 2014; Sanders *et al.*, 2015). Unfortunately, this number corresponds to only a small portion of the large genetic puzzle, which is expected to contain around a thousand genes (He *et al.*, 2013). Detection of *de novo* loss-of-function mutations (dnLoF) has been the key for gene discovery due to their high signal-to-noise ratio. However, such mutations are rare and they affect a diverse set of genes. Thus, for most of the genes, the rarity and diversity of variants prevent statistically significant signals from being observed. Therefore, our yield of discovered genes is still low, despite analyzing thousands of trios. The journey towards getting the full picture of the genetic architecture will take a long time and will be financially costly. Guilt-by-association based gene discovery techniques came handy to predict ASD risk genes. They assume these genes are working as a functional cluster. Starting from already known risk genes, these techniques predict a cluster of closely interacting genes. There are only a few network-based ASD-tailored gene discovery algorithms in the literature (Gilman *et al.*, 2011; Liu *et al.*, 2014; Hormozdiari *et al.*, 2014) that are later improved (Gilman *et al.*, 2012; Liu *et al.*, 2015) and these algorithms are described in the *Background* section. Despite their fundamental differences, all of these methods have one point in common: The biological gene interaction networks that they use are static. None considers the fact that gene interactions (coexpressions) evolve over time: It was clearly demonstrated that different neurodevelopmental spatio-temporal windows have different topologies and consequently, the clustering of ASD susceptibility genes drastically change (Willsey *et al.*, 2013). Moreover, dysregulation of pathways in earlier periods have cascading effects on the circuitry of the future time periods. For instance, it is shown that ASD risk increases ^36^-fold with a damage in early cerebellar circuitry, but a damage in adulthood does not cause dysfunction in social interactions (Wang *et al.*, 2014). Furthermore, early cerebellar damage has been shown to result in a cascade of long-term deficits in cerebellum and cause ASD (de la Torre-Ubieta *et al.*, 2016). Thus, we argue that the state-of-the-art methods are limited in their prediction power since static networks they use would fail to capture the dynamic nature of neurodevelopment.

In this article, we propose a novel ASD gene discovery algorithm termed ST-Steiner. The algorithm modifies the prize-collecting Steiner forest (PCSF) problem and extends it to spatio-temporal networks in order to mimic neurodevelopment. Instead of performing gene discovery on a single network or any number of networks separately, ST-Steiner solves an optimization problem progressively over a cascade of spatiotemporal networks while leveraging information coming from earlier neurodevelopmental periods. The algorithm has three novel aspects: (i) For the first time, the problem is solved on spatio-temporal coexpression networks and the dynamic nature of neurodevelopment is taken into account; (ii) The results are more interpretable compared to other methods in the literature, since the decisions made by the algorithm can be traced back on the spatio-temporal cascade; (iii) As the problem is formulated as a PCSF prediction problem, the algorithm predicts only the genes that are essential for connectivity of known risk genes. Note that, this is in contrast to the other approaches which can return redundant paths between known risk genes. We apply the algorithm on exome sequencing data from (De Rubeis *et al.*, 2014), and identify gene clusters using 2 gene coexpression network cascades of early-fetal and mid-fetal periods. For the first time, we benchmark the performances of the network-based ASD gene discovery algorithms on common training data. We validate the predicted clusters with independent exome sequencing data from (Iossifov *et al.*, 2014) and find that ST-Steiner’s predictions (i) identify more risk genes with less predictions compared to the state-of-the-art methods, (ii) overlap more with targets of known ASD-related transcription factors and enriched more in ASD-related KEGG pathways. We also show in various controlled settings that using temporal information boosts the prediction power, which supports our claim that the clustering of risk genes is spatio-temporal. We demonstrate that prediction of genes like NOTCH2 and NOTCH3 can be traced back to their role in neuron differentiation in embryonic period. Finally, we predict new genes with the incorporation of time dimension which point to a disruption of organization of microtubules in the embryonic period.

## II. BACKGROUND

### 1) Gene discovery in the context of neurodevelopmental disorders

Network-based gene discovery and disease gene prioritization have been very popular research areas in the last decade (Bebek *et al.*, 2012; Bromberg, 2013; Cowen *et al.*, 2017). While these methods proved their use mainly in the context of cancer, a parallel track has opened up for networkbased gene discovery for neurodevelopmental disorders, due to the specific characteristics of ASD and related disorders. The main reason is the rarity of dnLoFs and the small number of known seed genes. In addition, PPI networks do not suit well for the neuron specific analyses as they are mainly based on single isoform splicing (Corominas *et al.*, 2014; Hormozdiari *et al.*, 2014). There are only a few established network-based ASD gene discovery algorithms in the literature.

One among these methods, NETBAG, makes use of a gene interaction network which integrates numerous resources from the literature. Starting from the known ASD genes, NETBAG generates many clusters of genes at different sizes. Clusters are enlarged by adding the *closest* gene at every step in a greedy fashion. Each cluster is assigned a score that signifies the prior expectation (likelihood) that its genes contribute to the same genetic phenotype together. The algorithm returns a cluster with the maximal score. Then, using a permutation test, the significance of its score is established. NETBAG is initially designed to work with Copy Number Variations (CNV) data (Gilman *et al.*, 2011), but is extended to include other types of genetic variations (Gilman *et al.*, 2012).

A more recent method is DAWN, a hidden Markov random field based approach. It predicts ASD genes by assigning posterior risk scores to each gene, based on their interactions with other genes in a gene coexpression network (Liu *et al.*, 2014, 2015). Inputs to DAWN are prior TADA risk scores per gene (He *et al.*, 2013), and a partial coexpression network. A gene is assigned a higher risk score if the gene has a high prior risk, or is highly interacting with other risk genes.

Latest is the MAGI algorithm (Hormozdiari *et al.*, 2014). MAGI also uses a seed-and-extend based approach like NETBAG. Using a combination of PPI networks and a coexpression network, it generates seed pathways which are enriched with dnLoFs in cases, compared to controls. Then, it merges the pathways as long as the cluster score is improved. Finally, some genes are removed, added or swapped in order to reach a local maximum score. Using a permutation test, the significance of the detected modules are assessed. In addition, MAGI finds an extended module which is the union of high scoring suboptimal clusters that overlap with the best module. Besides, in order to obtain additional modules, genes of the best cluster are removed and MAGI is run again to obtain a new cluster.

A final method takes a different path: Even though it uses a brain specific functional gene interaction network, in contrast to ST-Steiner and the three methods mentioned above, it does not aim to predict a cluster of genes (Krishnan *et al.*, 2016). Instead, it uses the known ASD genes and their connectivity patterns as features to train an SVM classifier, which is used to assign every gene a probability of being associated to ASD.

### 2) No consensus on the network choice

Module detection methods in the literature assume that ASD risk genes must be clustered in a gene interaction network. Yet, the nature of the networks they prefer differs. NETBAG constructs its own network through a comprehensive analysis of many annotation resources (e.g. GO, protein domains), and numerous available interaction networks (e.g. PPIs, KEGG pathways). The method assigns each gene pair an interaction score that signifies the likelihood of those genes participating in a genetic phenotype. MAGI uses two PPIs: STRING (Szklarczyk *et al.*, 2010) and HPRD (Keshava Prasad *et al.*, 2008). It also makes use of the coexpression network generated by using full (all brain regions and neurodevelopmental periods) data from the BrainSpan dataset (Sunkin *et al.*, 2012). DAWN estimates partial coexpression networks using BrainSpan, but for only small windows of the neurodevelopment that are implicated as hotspots for ASD (Willsey *et al.*, 2013). Unlike NETBAG and MAGI, which use phenotype-neutral networks, DAWN’s approach favors links to already known ASD genes. A data-driven tissue-specific network is generated in (Krishnan *et al.*, 2016), where thousands of experiments from the literature are integrated in a Bayesian framework (Greene *et al.*, 2015). Despite such diverse design choices, all of the above-mentioned networks are static, i.e. their links do not change.

### 3) Steiner tree based formulations have proven to be useful in the biological domain

The method we propose to remedy the problems posed by using static networks is built on prize-collecting Steiner tree (PCST) problem. The original Steiner tree problem is concerned with finding a tree that connects a given set of distinguished (seed) nodes (Dreyfus and Wagner, 1971; Winter, 1987). It is first used over node-weighted biological networks to detect regulatory subnetworks (Scott *et al.*, 2005). The prize-collecting version of the problem aims to find a tree that maximizes the sum of the prizes of the selected nodes while penalizing the total cost of the connecting edges. In the biology domain, PCST has proven useful for applications like detecting functional modules in PPIs (Dittrich *et al.*, 2008), signaling pathway prediction (Huang and Fraenkel, 2009; Bailly-Bechet *et al.*, 2011), metabolic pathway prediction (Faust *et al.*, 2010) and drug repositioning (Sun *et al.*, 2016). Prize-collecting Steiner forest (PCSF) problem is a relaxation of PCST such that multiple disconnected components (trees) are allowed. PCSF has been used to identify multiple independent signaling pathways on a single network (Tuncbag *et al.*, 2013). Later, PCSF is extended to predict a single tree shared among multiple samples (networks with identical topology) with different mutation profiles (different seed genes) (Gitter *et al.*, 2014). The aim is to identify a common signaling pathway for diverse samples with the same phenotype.

### 4) Comparison with the literature

Among the state-of-the-art network-based ASD gene discovery methods, ST-Steiner is the first ASD gene discovery algorithm to consider the effect of the dysregulation in earlier periods of human brain development in its analysis. It uses a cascade of coexpression networks representing spatial and temporal windows of neurodevelopment. The algorithm solves the PCSF problem sequentially over this cascade of networks such that its selections on the target network (e.g. prefrontal cortex in midfetal period) is affected by the selections made in the earlier periods (e.g. prefrontal cortex in early fetal period). Unlike its counterparts, the reason a gene is selected can be traced back by following the decisions made in the earlier periods. Thus, the results are biologically more interpretable as the user gains insight into the flow of information from precursor time windows to the final predicted gene set.

Among the PCSF-variant approaches, we are only aware of two applications of PCSF on temporal data. The first one uses a temporal cascade like ours, however, tries to satisfy *k* connectivity demands (e.g. vertex1 must be connected to vertex2) (Khodaverdian *et al.*, 2016). This is different than our application, which aims to incorporate information from past time points without imposing such constraints. The second work uses fold changes in temporal expression data, but by solving PCSF on static PPI networks (Budak *et al.*, 2015). In ST-Steiner, selection of similar genes across multiple networks is rewarded in a similar manner to (Gitter *et al.*, 2014). However, in contrast, (i) ST-Steiner works with networks of different topologies, (ii) networks are organized in a temporal hierarchy, (iii) reward mechanism is weighted by the prize of a node and only affects networks of future time windows, and, networks represent spatio-temporal windows in brain development rather than samples with different mutation profiles. In addition, multiple brain regions in the same time window can be simultaneously analyzed without the constraint to be similar to each other. Then, all selected genes in this time frame can be assigned an additional prize in the next time frame analysis.

A motivating toy example for ST-Steiner and its decision making process is illustrated in Figure 1. On the coexpression network of spatio-time window 1, known risk genes (black) are connected by selecting (red-bordered) genes by solving PCSF. This result affects which genes will be selected on the coexpression network of spatio-temporal window 2. Assume genes *X, Y* and *Z* are equally likely to be selected (equal prizes and edge costs) to connect the seed genes. The algorithm prefers gene *X*, because it is selected in the earlier period and its prize is increased.

**Fig. 1:**
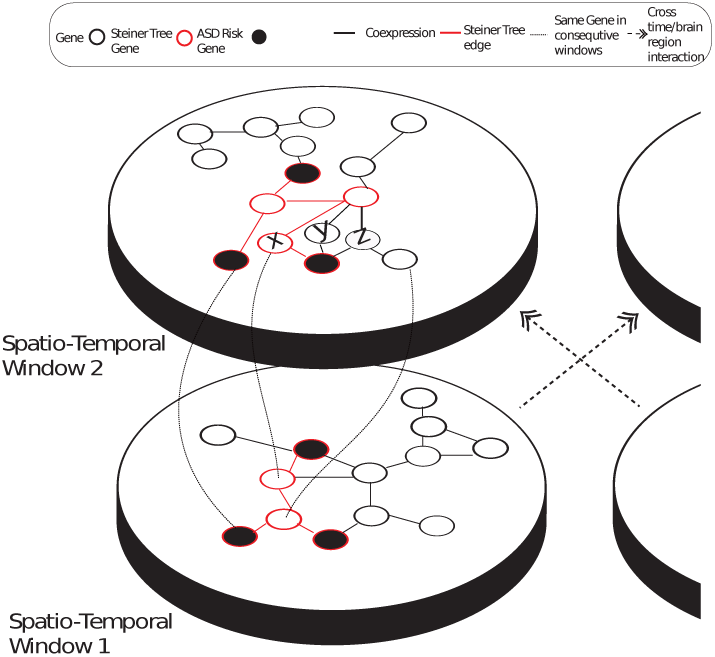
Figure shows 2 spatio-temporal windows (plates) and respective coexpression networks along with a parallel brain region and its plates (partially shown, on the right). Circles represent genes and black edges represent pairs of genes that are coexpressed. Red-bordered nodes form the Steiner tree found on plate 1 (linked with red edges), which minimally connects black seed genes. In ST-Steiner, genes that are selected in plate 1 are more likely to be selected in plate 2. Curved lines between windows show the mapping of selected genes from plate 1 to plate 2. On the second plate ST-Steiner can pick *X, Y* or *Z* to connect the seed genes. Assuming that they all have identical priors and identical edge costs, the algorithm would pick *X*, because it is selected in the prior window and its prize is increased. If other brain regions in the first temporal window are also considered, then selected genes in those regions would also be used (from the plate on the right).

## III. METHODS

### A. Prize-collecting Steiner tree problem (PCST)

Let *G*(*V, E*) denote an undirected vertex- and edge-weighted graph. *V* is the set of nodes, *E ⊆ V × V* is the set of edges that connects node pairs. *p*: *V →* ℝ_*≥*_0 is the node prize function and *c*: *E →* ℝ_*≥*_0 is the edge cost function for *G*. Given *G*, the task is to find a connected subgraph *T* (*V*_*T*_, *E*_*T*_) of the graph *G*, that minimizes the following objective function:

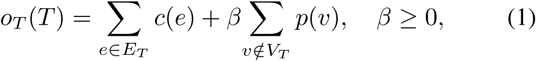

where *β* is a sparsity parameter that adjusts the trade-off between node prizes and edge costs. This trade-off corresponds to collecting the prize on a node by including the node and evading its edge cost by excluding it. An optimal solution is a tree, since if a cycle exists in *T*, any edge on the cycle can be removed to obtain another tree *T* ^*′*^ with *o*_*T*_ (*T* ^*′*^) *≤ o*_*T*_ (*T*).

### B. Prize-collecting Steiner forest problem (PCSF)

PCSF is an extension of PCST that lifts the connectedness constraint on the desired subgraph: Instead of a single tree, the goal is to find a forest. Given *G*, the problem is to find a subgraph *F* (*V*_*F*_, *E*_*F*_) of the graph *G* that minimizes the following objective function:

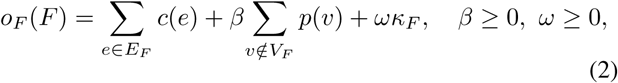

where *κ*_*F*_ *∈* N is the number of connected subgraphs (trees) in the subgraph *F* and *ω* is a parameter that adjusts its penalty. PCSF is a generalized version of PCST and reduces to PCST when *ω* = 0. An instance of PCSF can be solved as a PCST instance by adding an artificial node *v*0 to *V* and edges *E*0 = {*v*0*v*_*i*_ | *v*_*i*_ *V* with cost *ω* to *E*. Solving PCST on this new graph, and afterwards, removing *v*0 and *E*0, yields a minimal solution for the original PCSF instance (Tuncbag *et al.*, 2012).

### C. Spatio-temporal prize-collecting Steiner forest problem (ST-Steiner)

In order to model the spatio-temporal dynamics of neurodevelopment, we consider a spatio-temporal system ***G*** = (***G*_1_**, ***G*_2_**, *…,* ***G*_*T*_**), a list of *T* consecutive time windows.

The *i*^*th*^ time window 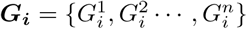 is a set of spatio-temporal networks, with a cardinality of *n*. The network 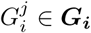(with node prize function 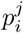 and edge cost function 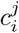), captures the topological state of the system ***G*** for the *j*^*th*^ spatial region in the *i*^*th*^ temporal window. In the context of spatio-temporal gene discovery for ASD, the network 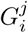 represents the coexpression of genes during human brain development at the *j*^*th*^ brain region cluster out of *n* regions in total during the *i*^*th*^ time interval [*t*_*i*_, *t*_*i*_ + *τ*], where *τ* ∈ N is the granularity parameter.

Given a spatio-temporal system ***G***, the problem is finding a minimum spatio-temporal sub-system ***F*** = (***F*_1_**, ***F*_2_**, …, ***F*_*T*_**). 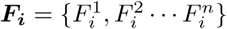 derives from the *i*^*th*^ time window ***G*_*i*_**, and 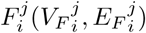 is a subgraph of graph 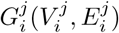. An optimal sub-system ***F*** minimizes the following objective function:

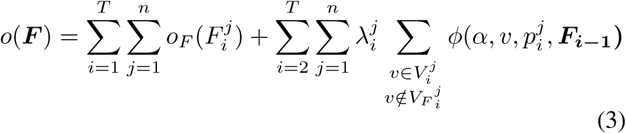

where (i) 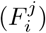 refers to the objective function for a single forest *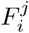*, shown in Equation 2, (ii) *ϕ* is an artificial prize function that promotes the selection of nodes which are selected in forests of the previous time window ***F*_*i-1*_**, and finally, (iii) 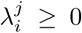 is a parameter that adjusts the impact of the artificial prize. The artificial prize function defined in Equation 4 is similar to the definition in (Gitter *et al.*, 2014), but here, each node gets an artificial prize proportional to its prize.

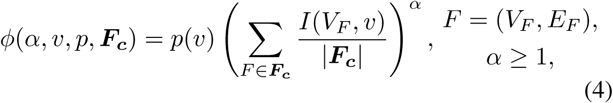

where (i) *α* adjusts the nonlinearity between the artificial prize for node *v* and the fraction of inclusion of node *v* among the set of subgraphs ***F*_*c*_**, and (ii) *I*(*V*_*F*_, *v*) is an indicator function that has value 1 if *v ∈ V*_*F*_, 0 otherwise. Note that, the use of function *ϕ* corresponds to increasing the prize 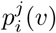 by an artificial prize, for all time windows *i >* 1 and each node 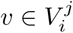, such that 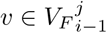 and *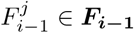*.

## IV. RESULTS

### A. Datasets and Generation of the Spatio-Temporal Networks

In order to model neurodevelopment, we use the BrainSpan microarray dataset of the Allen Brain Atlas (Sunkin *et al.*, 2012) and generate a spatio-temporal system (cascade) of coexpression networks. To partition the dataset into developmental periods and clusters of brain regions, we follow the practice in (Willsey *et al.*, 2013) as described next.

Brain regions are clustered according to their similarity as done in (Willsey *et al.*, 2013) and four clusters are obtained: (i) V1C (primary visual cortex and superior temporal cortex) (ii) PFC (prefrontal cortex and primary motor-somatosensory cortex), (iii) SHA (striatum, hippocampal anlage/hippocampus, and amygladia), and (iv) MDCBC (mediodorsal nucleus of the thalamus and cerebellar cortex). The time windows which are associated with these brain regions: 1–3 which corresponds to early fetal period, 3–5 and 4–6 which correspond to mid-fetal periods (*τ* = 2). Note that these time windows represent early neurodevelopment which is an important stage for ASD. Each graph 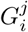 is a spatio-temporal coexpression network, where *i* denotes one of the time intervals and

*j* denotes one of the 4 brain region clusters. In this work, a spatiotemporal window of neurodevelopment and its corresponding coexpression network is denoted by the name of brain region cluster followed by the time-window of interest, e.g. PFC(1-3).

We report our results on the following two spatio-temporal cascades: (i) All spatial regions in time window 1–3, i.e. SHA(1-3), V1C(1-3), MDCBC(1-3) and PFC(1-3), as precursor networks for PFC(3-5) and (ii) all spatial regions in time window 1–3 as precursor networks for PFC(4-6). The target networks of interest, PFC(3-5) and PFC(4-6), are implicated as hotspots for ASD risk in (Willsey *et al.*, 2013). Furthermore, these are also the subject-matter of DAWN (Liu *et al.*, 2015), which allows us to directly compare our results to theirs.

An edge between two nodes is created if their absolute Pearson correlation coefficient *|r|* is *≥*0.7 in the related portion of BrainSpan data. This threshold has also been used in the literature (Liu *et al.*, 2014, 2015). Each edge between a pair of genes is assigned a cost of 1 *- r*2. We set node (gene) prizes to the negative natural logarithm of the TADA q-values. Thus, in all experiments, prize function is identical for all networks. TADA is a statistical framework that assigns each gene a prior risk by integrating information from *de novo* and inherited mutations as well as small deletions (He *et al.*, 2013; Sanders *et al.*, 2015). We obtain TADA q-values on ASC WES cohort which is reported on 17 sample sets consisting of 16,098 DNA samples 3,871 ASD cases (also 9,937 ancestry-matched/parental controls) from (De Rubeis *et al.*, 2014).

### B. Parameter Selection Procedure for ST-Steiner

The algorithm has 4 parameters for each spatio-temporal network. The first one is *ω*, which adjusts the number of trees in an estimated forest. This parameter is set to 0 to obtain a single tree per network, which is in line with the common assumption in the literature that ASD genes are part of a connected, functional cluster. *α* is the second parameter which adjusts the artificial prize and is set to 2 in all our tests to add modest non-linearity to the artificial prize, as done in (Gitter *et al.*, 2014). The third parameter is *β*, which adjusts the trade-off between the node prizes and edge costs. The final parameter is *λ*, which controls the magnitude of the artificial prize.

In order to select *β*, we make use of *efficiency ratio* (*ε*), which is defined as 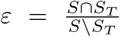, where *S*_*T*_ is a set of a ground truth (target) nodes to be covered by selecting a set of nodes *S* (Huang *et al.*, 2013). A high efficiency ratio is desirable, since it indicates that more target genes are covered by including less new predictions. In this work, we use 107 genes that are concluded as the predicted set of risk genes in (De Rubeis *et al.*, 2014) as our ground truth set *S*_*T*_. We experiment with two *ε* values and select *β* to yield *ε* = .5 *±t* or *ε* = .67 *± t*, with tolerance *t* = .05. Note that *ε* = .5 means that in *S*, there exist two additional genes that are not in *S*_*T*_ for each target gene in *S*_*T*_; similarly, *ε* = .67 means three extra genes are included in order to add two target genes. If no such network is found within a pre-specified number of runs (i.e. 10), the tolerance is relaxed to *t* = .1.

For each spatio-temporal network, to find a *β* that yields a desired level of *ε*, we first set *λ* to 0 (i.e. no artificial prize so the time dimension is ignored). Then, we perform a binary search for *β* on the fixed interval of [0.0, 2.0]. Note that, this procedure is performed for all coexpression networks of interest.

After fixing the *β*, using the same binary search procedure, we select a *λ* that promotes the addition of a small number of genes that are selected in the previous time-window. For this purpose, the selected subgraph *F*^*′*^(*V*_*F′*_ *E*_*F′*_) is targeted to have number of nodes *|V*_*F*′_ *|* = (1 + *ρ*) *|V*_*F*_*|± t*, where tree-growth parameter *ρ ≥* 0 targets a new size. We experiment with *ρ* = .05 and .1, which corresponds to growing the network in size by 5% and 10%, respectively. This overall procedure is performed consecutively in time-order for coexpression networks in all time-windows except the first time-window, where *λ* is set to 0.

### C. Comparison with the State-of-the-art Methods

We compare the performance of ST-Steiner with three state-of-the-art network-based ASD gene discovery algorithms which predict a module of ASD genes: NETBAG (Gilman *et al.*, 2011, 2012), DAWN (Liu *et al.*, 2014, 2015) and MAGI (Hormozdiari *et al.*, 2014).

#### 1) Input Training Data

The data from the ASC WES cohort from (De Rubeis *et al.*, 2014) is input to all three methods. ST-Steiner makes use of TADA values. NETBAG utilizes a list of genetic events: We treated each gene with one or more dnLoF as if it was hit by a separate event targeting that gene only. DAWN uses TADA values as in ST-Steiner. MAGI, by design, uses *de novo* counts.

#### 2) Input Networks

ST-Steiner uses the two cascades that are explained in the *Datasets and Generation of The Spatio-Temporal networks* subsection. For the other three methods, the suggested networks in the corresponding papers are used. That is, NETBAG is run with its own phenotype network. DAWN uses the PFC(3-5) and PFC(4-6) partial coexpression networks obtained from BrainSpan as reported in (Liu *et al.*, 2015). MAGI is run using the STRING (Szklarczyk *et al.*, 2010) and HPRD (Keshava Prasad *et al.*, 2008) PPI networks and the full coexpression network obtained from the BrainSpan dataset (Sunkin *et al.*, 2012).

#### 3) Input Parameters and Implementation Details

ST-Steiner parameters are selected as described in the *Parameter Selection for ST-Steiner* subsection. Here, for both cascades, we report results obtained by using *ε* = .5 and *ρ* = .1 unless stated otherwise. The ST-Steiner results obtained by using different parameters (other than *ε* = .5 and *ρ* = .1) are stated explicitly while denoting the result. The exact parameter values for *β, λ, ε* and *ρ* for every ST-Steiner result in this manuscript are reported in Supplementary Tables 1–3. We use a message-passing based implementation named msgsteiner(v1.3)^1^ to solve PCST (Bailly-Bechet *et al.*, 2011). We fix the solver’s parameters as follows: The maximum search depth to 30, noise to 0 and the reinforcement parameter to 1*e -* 3. The rest are set to their defaults.

**TABLE I:** Overlap of predicted gene sets of each method with the 251 genes that have at least 1 dnLoF in 1,643 samples from (Iossifov *et al.*, 2014) which are not included in the training set. Fisher’s exact test is used to assess significance. The most significant result is marked in **bold**. As for the background 18,736 genes are used (the union with TADA genes in (De Rubeis *et al.*, 2014) and an extra gene included by MAGI). Note that MAGI Best1 *⊂* MAGI Ext1, but MAGI Best2 */⊂* MAGI Ext2.

| gene set name | p-value | intersection / # predictions |
| --- | --- | --- |
| ST-St. Every(1-3)+PFC(3-5) | **6.958e-12** | **21 / 234** |
| ST-St. Every(1-3)+PFC(4-6) | 1.827e-09 | 19 / 256 |
| NETBAG | 2.476e-06 | 9 / 87 |
| DAWN PFC(3-5) | 3.842e-08 | 17 / 246 |
| DAWN PFC(4-6) | 9.017e-10 | 18 / 218 |
| MAGI Best1 | 3.679e-05 | 6 / 47 |
| MAGI Ext1 | 7.865e-05 | 8 / 104 |
| MAGI Best2 | 2.628e-02 | 2 / 19 |
| MAGI Ext2 | 4.396e-03 | 5 / 80 |

NETBAG is run with its latest available code and data, dated January 2016^2^. We follow the steps presented in the README file provided with the code package. We prepared a genetic events file that lists every gene with at least one dnLoF mutation observed in the same input WES data, as separate events. We performed 10,000 null distribution trials, as in the original paper (Gilman *et al.*, 2011). For the gene metrics data required by the algorithm, we supplied the data file that contains the average connectivity of the top 20 genes, which is included in the NETBAG code package. For DAWN, since the reported gene sets in (Liu *et al.*, 2015) are also generated with the same input WES data, we use the lists as they are. MAGI is run with its publicly available code (v0.1 Alpha^3^). We set the upper bound on the size of a module to 200. For the rest of the required parameters, values from the paper are used (Hormozdiari *et al.*, 2014). Optional parameters are set to their default values as supplied in their manual file. We ran MAGI for 310,000+ iterations to obtain the first set of predictions (M1 Best and M1 Extended). After M1 Best is removed, MAGI is run for another 310,000+ iterations to obtain M2 Best and M2 Extended —see (Hormozdiari *et al.*, 2014) for details.

#### 4) Resulting predictions

ST-Steiner predicts a network of 234 genes on the first cascade: This resulting network is referred to as ST-St. Every(1-3)+PFC(3-5). On the second cascade, it predicts a network of 256 genes and it is referred to as ST-St. Every(1-3)+PFC(4-6). See the gene lists in Supplementary Tables 4 and 5, respectively.

NETBAG selects a cluster of 87 genes, with global p-value 0.048, local p-value 0.009, z-score 0.2209 and cluster score 139.486. The list of genes is given in Supplementary Table 6. DAWN outputs two networks provided in (Liu *et al.*, 2015): (i) DAWN PFC(3-5) (246 genes), (ii) DAWN PFC(4-6) (218 genes). MAGI predicts a core network with 47 genes (M1 Best —referred to as MAGI Best1), which is then extended to 104 genes (M1 Extended —referred to as MAGI Ext1). Second core network (M2 Best —referred to as MAGI Best2) contains 19 genes which is then extended to module M2 Extended (MAGI Ext2, 80 genes). These gene lists are provided in Supplementary Tables 7 and 8. Overlaps between the predictions of ST-Steiner, NETBAG, DAWN and MAGI are given in Supplementary Table 9.

#### 5) Validation of the Predicted Networks on Independent WES Data

In this experiment, we validate the predicted gene networks by each method, using the dnLoFs observed in 1,643 samples in the SSC WES cohort which are not included in the input training data (which is ASC WES cohort). A dnLoF mutation has a very high signal-to-noise ratio, and the genes that are hit are considered to be high risk genes. Therefore, since the two cohorts are completely independent, this is a powerful validation experiment and constitutes a benchmark for all methods, as also done in (Liu *et al.*, 2015). Note that we are also comparing all methods when they are provided with the identical training data (of ASC WES cohort): In this sense, this work is the first benchmark of all network-based ASD gene discovery algorithms in the literature when the training data is the same. There are 251 such genes (that are hit in the SSC cohort) and we call them validated genes from this point on. Inspecting Table I for the results of this test, we see that ST-St. Every(1-3)+PFC(3-5) returns the most significant overlap by recovering 21 of those genes with only 234 predictions. DAWN PFC(4-6) gets the second most significant result by identifying 18 validated genes with 218 predictions and ST-St. Every(1-3)+PFC(4-6) is ranked third with 19 validated genes but with 256 predictions. On the other hand, MAGI and NETBAG are not as successful as ST-Steiner and DAWN in identifying these genes.

One important observation is that despite using the same spatio-temporal windows and having 56.8% and 67.4% overlaps with the DAWN PFC(3-5) and DAWN PFC(4-6) networks, ST-Steiner’s predictions recover more validated genes. This can be attributed to both (i) using the cascaded information coming from a preceding neurodevelopmental window and (ii) ST-Steiner predicting a tree (rather than a forest), which only includes high prize low cost genes that are essential for connectivity. We validate the benefit of utilizing the time dimension further in the *Alternative Uses of Spatial and Temporal Information* subsection.

Aside from the state-of-the-art methods described above, we also compare the same predictions of ST-Steiner with other predicted ASD-risk gene modules from the literature: NETBAG (Gilman *et al.*, 2011), AXAS (Cristino *et al.*, 2014), and coexpression based modules from (Willsey *et al.*, 2013) and (Parikshak *et al.*, 2013). As done in (Hormozdiari *et al.*, 2014), we take their outputs as is for these comparisons. Results are given in Supplementary Table 10. We note that none of the predicted modules get close to the precision of ST-Steiner.

#### 6) Validation of the Predicted Networks on ASD-related Gene Sets Indicated in Other Resources

While the exact set of functional circuits and a full picture for the genetic architecture of ASD is far from complete, there are some leads: We compare the overlap of predicted gene sets by each method with gene sets that are indicated as relevant to ASD in external resources. Fisher’s exact test is used to assess significance. We present the results in Table II and indicate the most significant overlap values in bold.

First, we compare the overlaps of the results of each method with the Simons Foundation Autism Research Initiative - SFARI’s Category 1 and 2 genes^4^. Category 1 includes 24 ASD-associated genes with high confidence that are confirmed by independent experiments. Category 2 consists of 59 genes with strong confidence —without requiring independent replication of association. ST-St. Every(1-3)+PFC(3-5) predicts the largest number of Category 1 genes (16 out of 24), while DAWN PFC(4-6) predicts more genes in Category 2 (21 out of 59).

Next, we consider the targets of two genes which are previously shown to be related to ASD. FMRP is a transcription factor whose targets have been shown to be associated with ASD (Iossifov *et al.*, 2012). These targets are proposed to be used as extra covariates in DAWN algorithm and are found to be significantly contributing to the prediction performance (Liu *et al.*, 2015). Furthermore, the identified ASD-risk genes have shown significant overlaps with these targets (De Rubeis *et al.*, 2014). RBFOX, on the other hand, is a splicing factor and was shown to be dysregulated in ASD (Weyn-Vanhentenryck *et al.*, 2014; Voineagu *et al.*, 2011). Its targets also have been shown to overlap with ASD genes (De Rubeis *et al.*, 2014). We compare the overlaps between the results of each method and (i) 842 FMRP targets reported in (Darnell *et al.*, 2011), 1,048 RBFOX targets reported in (Weyn-Vanhentenryck *et al.*, 2014) with significant HITS-CLIP peaks, and finally 587 genes which are RBFOX targets with alternative splicing events (Weyn-Vanhentenryck *et al.*, 2014; Voineagu *et al.*, 2011; De Rubeis *et al.*, 2014). For all three downstream target sets, ST-Steiner’s predictions have the most significant overlaps compared to the sets predicted by NETBAG, DAWN and MAGI. ST-St. Every(1-3)+PFC(4-6) identifies 25% more FMRP targets than the closest result. This finding stands out in these set of comparisons as the most significant overlap.

Then, we focus on two among the most well-established disrupted functionalities in ASD, which are also the main theme of (De Rubeis *et al.*, 2014) in addition to shaping its title: In ASD, the genes related to chromatin modification and synapses are disrupted. We obtain the list of 152 histone modifier genes (HMG) also used in (De Rubeis *et al.*, 2014) for enrichment analyses. ST-St. Every(1-3)+PFC(3-5) identifies 12 HMGs with 234 predictions and provides the most significant result. DAWN and MAGI return a maximum of 9 and 8 HMGs, respectively.

Finally, we construct a large set of synaptic genes by getting the union of synapse-related GO-terms —see the caption of Table II for the list of terms. MAGI has the most significant overlap in this category, by identifying 9 genes with only 19 predictions on the core of the second module —MAGI Best2. Although ST-St. Every(1-3)+PFC(4-6) identifies 27 synaptic genes with 256 predictions, it is not as significant due to its larger gene set size.

Comparison of ST-Steiner with other predicted clusters from the literature is shown in Supplementary Table 11 (Gilman *et al.*, 2011; Willsey *et al.*, 2013; Cristino *et al.*, 2014; Parikshak *et al.*, 2013). Out of many modules, M1 and M2 of (Cristino *et al.*, 2014) and M16 of (Parikshak *et al.*, 2013) show higher enrichment in HMGs, Synaptic genes and FMRP/RBFOX targets, respectively. However, these are very large sets (containing 700, 680 and 492 genes, respectively) and show very little overlap with known or likely ASD genes: SFARI Category 1 (0, 4 and 1 overlaps, respectively), SFARI Category 2 (9, 8 and 3 overlaps) and validated genes from (Iossifov *et al.*, 2014) (13, 7 and 7 overlaps).

#### 7) KEGG Pathway and GO Term Enrichment Comparison

We compare the performance of the methods with respect to their enrichment in KEGG pathways and GO terms. In order to generate lists of ASD-related KEGG pathways and GO Biological Process (BP) terms, we obtain the pathways and terms that significantly overlap with SFARI Category 1 and 2 genes (83 genes) using Enrichr tool with its gene-set libraries KEGG 2016 and GO Biological Process 2017 (Chen *et al.*, 2013). We obtain 6 such KEGG Pathways and 15 such GO BPs. Using Fisher’s exact test, we assess the significance of the overlap of the predictions of each method (ST-Steiner, NETBAG, DAWN and MAGI) with these pathways and terms. Results indicate that ST-Steiner has the most significant overlap in 3 out of 6 associated KEGG pathways and in 5 out of 15 associated GO terms, whereas MAGI outweighs in only 1 KEGG pathway and in 7 GO terms —see Supplementary Figures 2 and 3.

**Fig. 2:**
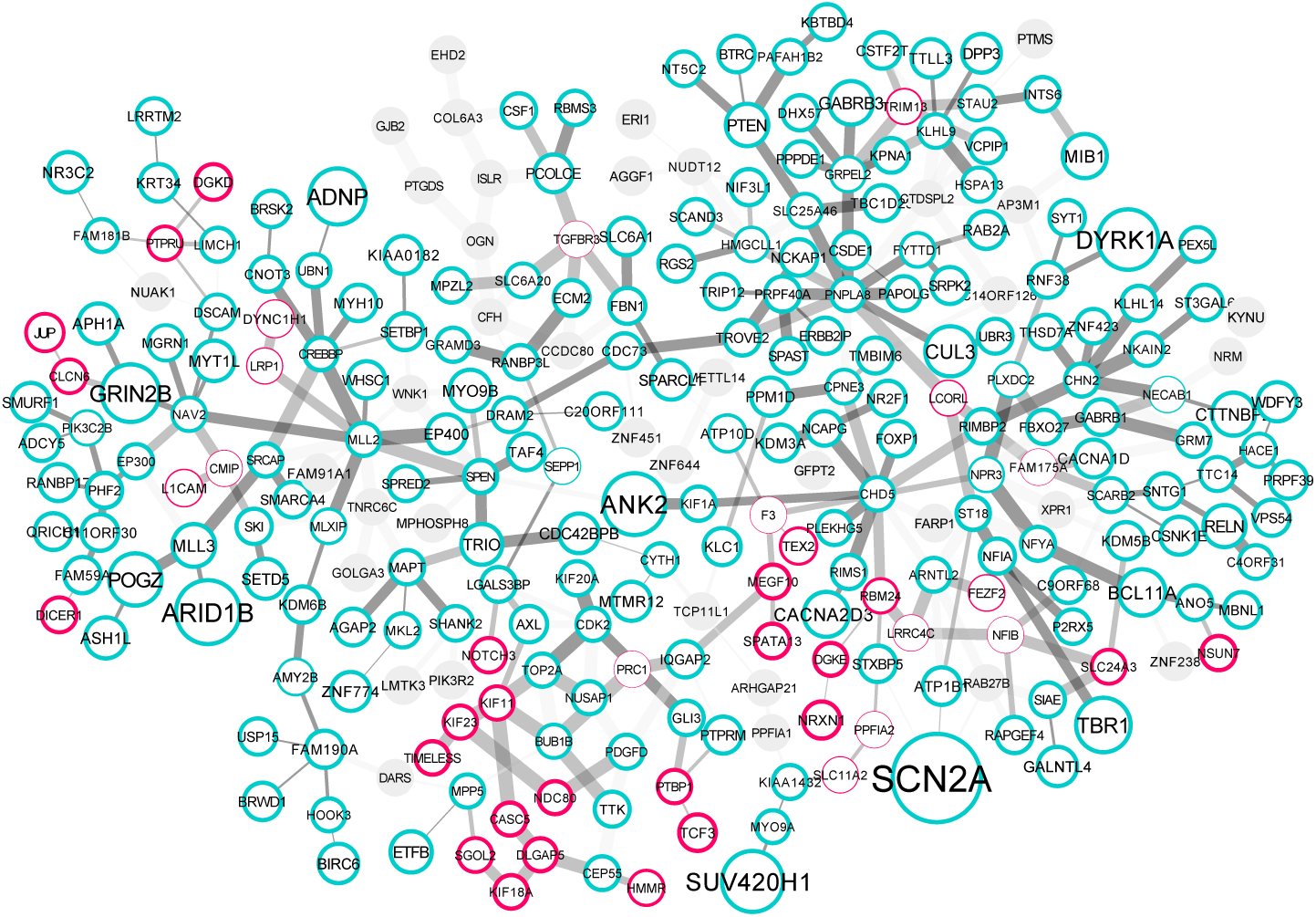
Figure visualizes ST-St. PFC(1-3)+(3-5) network which is laid over ST-St. PFC(3-5)+10%. Turquoise bordered genes are common in ST-St. PFC(1-3)+(3-5) and ST-St. PFC(3-5)+10%. Pink bordered genes are only in ST-St. PFC(1-3)+(3-5), highlighting the effect of using the temporal dimension and information coming from PFC(1-3). Gray genes are present only in PFC(3-5)+10% and are not included by ST-St. PFC(1-3)+(3-5). Size of a node indicates its significance w.r.t. its TADA q-value in (De Rubeis *et al.*, 2014) (the larger, the more significant). The thickness of the border of a node indicates its robustness. The thickness of an edges represents the correlation coefficient between the gene pair (the thicker, the higher). Visualized using the CoSE layout (Dogrusoz *et al.*, 2009) in Cytoscape (Shannon *et al.*, 2003).

As done in (Hormozdiari *et al.*, 2014), we obtain the top-3 pathways and GO BP terms for each method and compare every method on the union of these pathways and terms. Three notable findings related to ASD in these figures are (i) *GABAergic Synapse* term (ST-Steiner is better), (ii) *WNT Signaling pathway* (MAGI is better) and (iii) *Notch Signaling Pathway* (MAGI is slightly better than ST-Steiner) —see Supplementary Figures 4 and 5.

MAGI’s GO BP enrichment performance is better compared to other methods. ST-Steiner picks more relevant genes in related KEGG pathways. In this sense, MAGI and ST-Steiner perform comparably.

### D. Alternative Uses of Spatial and Temporal Information

In this section, we compare the results of ST-Steiner under different settings by discussing how the parameters and number of precursor windows affect the performance. Then, we evaluate the benefit of using temporal hierarchy. We use the same input data and parameters as described in the *Results* section. We consider two new cascades: (i) ST-St. PFC(1-3)+(3-5), where PFC(1-3) is the precursor window for PFC(3-5) and (ii) ST-St. PFC(1-3)+(4-6) —see Supplementary Tables 12 and 13 for detailed results on all upcoming comparisons in this section, and Supplementary Tables 14–19 for the referenced ST-Steiner results.

**TABLE II:** Table shows the significance of the intersection (*∩*) of the gene set *S* predicted by each method and ASD-related gene sets as indicated in external resources. Significance is assessed using Fisher’s exact test. As for the background 18,736 genes are used (the union genes with TADA values in (De Rubeis *et al.*, 2014) and 1 extra gene included by MAGI). The most significant result is marked in **bold**. The first two lists are SFARI Gene’s Category 1 (24 genes) and Category 2 (59 genes) gene sets. The third list is the list of transcription factor FMRP’s targets (842 genes). Fourth and the fifth gene sets are the targets of the splicing factor RBFOX: *RBFOX-peak* denotes 1,048 target genes with significant HITS-CLIP peaks and *RBFOX-splicing* denotes 587 genes RBFOX targets with alternative splicing events. Histone modifiers (152 genes) is the set of histone modifier genes used in (De Rubeis *et al.*, 2014). Finally, Synaptic genes (878 genes) is the union of synapse-related-GO terms’ gene sets which are GO:0021681, 0021680, 0021694, 0021692, 0060076, 0060077, 0051965, 0090129, 0032230, 0051968, 0045202, 0098794, 0098793, 0048169, 0051963, 0090128, 0032225, 0032228, 0051966, 0007271, 0001963, 0035249. Note that MAGI Best1 *⊂* MAGI Ext1, but MAGI Best2 *⊄* MAGI Ext2.

| Predicted Gene Set $S$ | SFARI Category 1 | | SFARI Category 2 | | FMRP Targets | | RBFOX - peak | | RBFOX - splice | | Histone Modifiers | | Synaptic Genes | |
| --- | --- | --- | --- | --- | --- | --- | --- | --- | --- | --- | --- | --- | --- | --- |
| | p-value | $\vert \cap\vert /\vert S\vert $ | p-value | $\vert \cap\vert /\vert S\vert $ | p-value | $\vert \cap\vert /\vert S\vert $ | p-value | $\vert \cap\vert /\vert S\vert $ | p-value | $\vert \cap\vert /\vert S\vert $ | p-value | $\vert \cap\vert /\vert S\vert $ | p-value | $\vert \cap\vert /\vert S\vert $ |
| ST-St. Every(1-3)+PFC(3-5) | **1.397e-25** | **16 / 234** | 2.906e-22 | 19 / 234 | 3.198e-22 | 52 / 234 | 2.196e-09 | 38 / 234 | **1.120e-11** | **31 / 234** | **4.709e-07** | **12 / 234** | 1.195e-04 | 25 / 234 |
| ST-St. Every(1-3)+PFC(4-6) | 8.334e-23 | 15 / 256 | 5.890e-20 | 18 / 256 | **8.684e-27** | **60 / 256** | **1.735e-11** | **44 / 256** | 3.649e-08 | 27 / 256 | 7.890e-06 | 11 / 256 | 7.622e-05 | 27 / 256 |
| NETBAG | 5.062e-18 | 10 / 87 | 1.422e-13 | 10 / 87 | 1.750e-10 | 21 / 87 | 2.870e-04 | 14 / 87 | 8.199e-05 | 11 / 87 | 7.103e-04 | 5 / 87 | 2.409e-03 | 11 / 87 |
| DAWN PFC(3-5) | 5.479e-19 | 13 / 246 | **4.242e-25** | **21 / 246** | 3.985e-18 | 48 / 246 | 2.740e-09 | 39 / 246 | 3.774e-09 | 28 / 246 | 1.835e-04 | 9 / 246 | 3.274e-04 | 25 / 246 |
| DAWN PFC(4-6) | 7.067e-24 | 15 / 218 | 1.252e-19 | 17 / 218 | 7.762e-19 | 46 / 218 | 2.735e-10 | 38 / 218 | 2.231e-08 | 25 / 218 | 7.347e-05 | 9 / 218 | 1.744e-03 | 21 / 218 |
| MAGI Best1 | 3.554e-07 | 4 / 47 | 9.615e-03 | 2 / 47 | 1.031e-08 | 14 / 47 | 3.943e-03 | 8 / 47 | 5.946e-04 | 7 / 47 | 2.100e-06 | 6 / 47 | 2.813e-01 | 4 / 47 |
| MAGI Ext1 | 1.866e-07 | 5 / 104 | 3.220e-04 | 4 / 104 | 3.600e-06 | 17 / 104 | 1.500e-02 | 12 / 104 | 2.137e-05 | 13 / 104 | 2.072e-06 | 8 / 104 | 9.394e-03 | 11 / 104 |
| MAGI Best2 | 2.652e-04 | 2 / 19 | 1.000e+00 | 0 / 19 | 2.321e-12 | 12 / 19 | 3.127e-03 | 5 / 19 | 2.512e-03 | 4 / 19 | 1.000e+00 | 0 / 19 | **6.258e-08** | **9 / 19** |
| MAGI Ext2 | 3.064e-06 | 4 / 80 | 2.049e-03 | 3 / 80 | 1.165e-08 | 18 / 80 | 1.442e-03 | 12 / 80 | 3.735e-05 | 11 / 80 | 1.000e+00 | 0 / 80 | 3.862e-06 | 15 / 80 |

#### 1) Effect of Parameters

First, we run ST-Steiner by setting *ρ* to .05, which decreases the number of time induced genes. Secondly, we increase *ε* to .67, which shrinks the size of the network. In both cases, the number of validated genes that are identified do not improve. Also, the number of genes predicted in our chosen setting is comparable with the literature (in particular, with DAWN). Hence, we fix our setting to *ε* = .5 and *ρ* = .1.

#### 2) Effect of Using Multiple Spatial Regions

We assess the effect of changing the number of spatial (brain) regions used as precursors. Results show that using a single precursor coexpression network rather than multiple reduces the overlaps with the lists from external sources. Therefore, we prefer using multiple precursors.

#### 3) ST-Steiner vs. Union of Independent Results

One way of joining information in two temporal windows is to analyze them independently and then, take the union of the selected genes. Yet, this would not predict a connected network of genes, which defies the initial assumption that the genes are part of a functional cluster. Still, we run ST-Steiner on PFC(1-3), PFC(3-5) and PFC(4-6) independently, with no time effect (*λ* = 0, *ρ* = 0, *β* is selected such that *ε* = .5) and then, obtain ST-St. PFC(1-3) *∪* ST-St. PFC(4-6). This results in a gene set of 293 genes, which does not predict any new validated genes, when compared to ST-St. PFC(1-3)+(4-6). *∪* ST-St. PFC(1-3) ST-St. PFC(3-5) has one more validated gene retained in the union set which includes 275 genes. However, due to its large size, it not as significant as ST-St. PFC(1-3)+(3-5) (*p* = 1.472*e -* 10 vs. 7.0*e -*12). Thus, our scheme is better in terms of the prediction performance and also for returning biologically interpretable results as the gene set is connected.

#### 4) ST-Steiner vs. Single Analysis with Coarser Granularity

To combine information in two windows, another alternative is to perform the analysis on a coexpression network that spans both windows (i.e. instead of PFC(1-3) and PFC(4-6), use PFC(1-6)). To assess this approach, we obtain network ST-St. PFC(1-6) with no time effect (*λ* = 0, *ρ* = 0, *β* is selected such that *ε* = .5). This network contains 197 genes and identifies only 14 genes that are validated (*p* = 4.480*e -*07). In contrast, our stepwise approach ST-St. PFC(1-3)+(4-6) identifies 19 validated genes by predicting 256 genes (*p* = 1.836*e-* 9). Similarly, ST-St. PFC(1-5) with no time effect also predicts only 14 validated genes and ST-St. PFC(1-3)+(3-5) identifies 21 validated genes. These indicate that finer granularity coexpression networks are more potent.

#### 5) ST-Steiner vs. No Temporal Effect

In order to see if information coming from a precursor improves the performance, we compare ST-St. PFC(1-3)+(3-5) and ST-St. PFC(1-3)+(4-6) with ST-Steiner results only on PFC(3-5) and PFC(4-6), respectively. For this purpose, we obtain ST-St. PFC(3-5) and ST-St. PFC(4-6) independently, with no time effect (*λ* = 0, *ρ* = 0, *β* is selected such that *ε* = .5). The former identifies 20 validated genes by making 211 predictions (*p* = 8.272*e -*12), while ST-St. PFC(1-3)+(3-5) identifies 21 validated genes by choosing 234 genes (*p* = 7.0*e -* 12). PFC(4-6) identifies 18 validated genes by predicting 235 genes (*p* = 3.018*e -* 09), whereas ST-St. PFC(1-3)+(4-6) identifies 19 validated genes by predicting 256 (*p* = 1.836*e -* 9). The increase in the number of validated genes demonstrate that ST-Steiner can leverage temporal information.

Due to *ρ*, the network size is increased to include more genes by considering the time dimension. Therefore, the above comparison is slightly unfair. What we want is to see if the leveraged information comes just from network getting bigger or is due to the spatio-temporal analysis. To investigate this, we obtain two additional results: ST-St. PFC(3-5)+10% and ST-St. PFC(4-6)+10% with the following parameters: *λ* = 0, *β* is selected such that they have comparable sizes to PFC(1-3)+(3-5) and PFC(1-3)+(4-6), respectively. These results contain 230 and 266 genes, respectively. We see that PFC(3-5)+10% and ST-St. PFC(4-6)+10% also could identify 20 and 19 genes respectively, which are less significant. Hence, we conclude that the prize mechanism successfully promotes selection of ASD-risk genes.

We also perform a robustness analysis similar to the one in Liu *et al.* 2014. That is, we remove the genetic signal from 1/30 randomly selected genes and rerun ST-Steiner. This is repeated 30 times for each fold to see how frequently each gene is selected. The visualization of ST-St. PFC(1-3)+(3-5) in comparison to ST-St. PFC(3-5)+10% is illustrated in Figure 2 along with the robustness values. See Supplementary Figure 1 for a similar comparison of ST-St. PFC(1-3)+(4-6) and PFC(4-6)+10%.

### E. Time and Memory Performance

All tests are performed on a server with 2 CPUs and 20 cores in total (Intel Xeon Processor E5-2650 v3) with 256GB memory. ST-Steiner’s running time scales with the size and connectivity of the input network as well as the number of networks considered in the cascade. The largest network we experiment with is PFC(1-3), which has 11,891 genes and 4,987,350 connections: ST-Steiner took *∼*6 hours with peak memory usage of 12 GB on maximum 40 threads.

## V. DISCUSSION

### 1) ST-Steiner predicts new genes with temporal information

We evaluate genes that are both included in ST-St. PFC(1-3)+(3-5) and ST-St. PFC(1-3)+(4-6), but not included in PFC(3-5)+10% or PFC(4-6)+10%, i.e. nodes that are predicted due to temporal information. There are 9 such genes and we focus on 6 which are detected neither by DAWN nor by MAGI. Furthermore, these genes are not pointed out in WES studies. They all have low scores in TADA, and ranked at most as Category 4 in SFARI Gene, if included at all.

The first one is KIF23, a kinesin family member protein. Kinesins are known to transport cargo to dendritic spines undergoing synaptic plasticity over microtubules (McVicker *et al.*, 2016). ST-St. PFC(1-3)+(3-5) selects 5 KIF genes. Note that TADA q-values are available only for 6 KIF genes. Kinesins are also known to play a role in organization of spindle microtubules during mitosis (Ems-McClung and Walczak, 2010). We detect two more genes, NDC80 and SGOL2, which take part in kinetochore-microtubule attachment during cell division. These two genes and above-mentioned KIF genes closely interact in Figure 2. We do not find any evidence of association with ASD for these 3 genes. These results suggest that an early disruption of these circuitries can be related to ASD.

The following genes have subtle clues in the literature. The fourth gene MEGF10 is also related to cell proliferation and adhesion, and recently variants in its regulatory region have tied it to ASD (Wu *et al.*, 2017). The fifth gene is CMIP, which is a signaling protein previously associated with language delay. There are two studies (one very recent) indicating haplosufficiency of CMIP leads to ASD (Van der Aa *et al.*, 2012; Luo *et al.*, 2017). Selection of CMIP in ST-St. PFC(1-3)+(3-5) enables inclusion of another gene L1CAM with a link to CMIP, which is related to neurite outgrowth and cell migration. The final gene, which is time induced (ironically), is TIMELESS. It is related to circadian rhythm and also linked to ASD before in a single study (Yang *et al.*, 2016).

### 2) Predicted genes provide traceable information

Here, we focus on genes that are relatively more established in literature compared to the genes in the previous subsection. However, them not being selected by ST-Steiner without the time information allows us to traceback the information and show that ST-Steiner is able to also return more interpretable results.

The first two genes are NOTCH2 and NOTCH3, membrane receptors of NOTCH signaling pathway. The former gene is selected in both ST-St. PFC(1-3)+(3-5) and ST-St. PFC(1-3)+(4-6), the latter is selected in ST-St. PFC(1-3)+(4-6). This pathway is important for neuron differentiation (Cau and Blader, 2009). More importantly, it is active during embryogenesis (time-point 1) (Wolter, 2013). Thus, its disruption is expected to have a cascaded effect during time-window 3–5. The second gene that is retained is TCF3, which has been highlighted in Figure 4 of (De Rubeis *et al.*, 2014) as one of the few hub transcription factors to regulate many ASD-risk genes along with many histone modifiers. Similar to NOTCH signaling pathway, which regulates neuron differentiation, TCF3 is found to promote differentiation in embryonic stem cells (Merrill *et al.*, 2001; Nguyen *et al.*, 2006). More importantly, it represses neuron differentiation in neural precursor cells (Kuwahara *et al.*, 2014; Ohtsuka *et al.*, 2011), which again corresponds to the time window 1–3. Thus, in line with our hypothesis that clustering of genes is spatio-temporal, ST-Steiner predicts these genes by considering the effect of the earlier time-window. This also means that one can trace back the information source which enables the selection of these genes: This adds further interpretability to our results.

## VI. CONCLUSION

ASD is a common neurodevelopmental disorder which is a life-long challenge for many families all around the world. Gaining an understanding of the cause of ASD and opening a way to the development of new treatments would certainly have a major impact on the lives of many. Even though we have long ways to go to understand the etiology of the disorder, network-based gene discovery algorithms have proven useful for gene discovery. In this work, we propose a novel ASD gene discovery algorithm that for the first time, models the cascading effect of disrupted functional circuits in neurodevelopment.

We show that ST-Steiner is more precise than the state-of-the-art methods. While other methods output a graph of genes, ST-Steiner outputs a tree/forest. That is, no multiple paths exist between any pair of genes and some genes can be left out if they are not essential for connectivity due to having low prior risk or high-cost edges. Precision is also achieved via rewarding selected genes from earlier periods, which makes the algorithm more confident on the predictions it makes. We show that once the temporal information is employed, prediction power increases, which supports our hypothesis that the clustering of genes is spatio-temporal rather than static.

We restrict our tests to ASD and only to well known ASD-related spatio-temporal windows (PFC(3-5) and PFC(4-6)) for a comprehensive comparison with other methods. However, ST-Steiner can be used for any disorder given a progression model and corresponding cascade of coexpression networks. One example is intellectual disability (ID). As a matter of fact, ID and ASD are known to be highly comorbid and to share a genetic component (McCarthy, 2007). It is an interesting problem to identify periods of divergence and convergence using ST-Steiner. One other target could be neurodegenerative disorders such as Alzheimer’s for which adulthood Brain coexpression networks from BrainSpan can be used.

We also think there is still room for improvement in ST-Steiner’s formulation. Considering that PPI networks give MAGI an orthogonal source of information, one future extension direction would be the incorporation of information from this source. Note that the goal of this paper is to show that the clustering of ASD genes are dynamic rather than static. This is the reason why we do not consider PPIs, but restrict the problem definition and our tests to observe the cascaded effects of dysregulation in earlier time windows.

## VII. AVAILABILITY

ST-Steiner code and a toy example are available on ST-Steiner website (http://ciceklab.cs.bilkent.edu.tr/st-steiner) under the GNU General Public License 3.0.

## VIII. ACKNOWLEDGEMENTS

We thank Fereydoun Hormozdiari and Li Liu for help with their methods, Oznur Tastan and Serhan Yilmaz for their feedback and Lambertus Klei for the help on processing BrainSpan data.

*1) Conflict of interest statement:* None declared.

## Footnotes

1 http://staff.polito.it/alfredo.braunstein/code/

2 http://netbag.org/

3 https://eichlerlab.gs.washington.edu/MAGI/

4 https://gene.sfari.org/database/gene-scoring accessed on Jan 15, 18

